# Spatiotemporal dynamics of questing activity by four tick species in the central Great Plains

**DOI:** 10.1101/2025.07.15.664906

**Authors:** Marlon E. Cobos, Joanna L. Corimanya, Eric Ng’eno, Claudia Nuñez-Penichet, Abigail C. Perkins, Zenia Ruiz-Utrilla, Abdelghafar Alkishe, Daniel Romero-Alvarez, Yuan Yao, Anuradha Ghosh, Xiangming Xiao, A. Townsend Peterson, Kathryn T. Duncan

**Affiliations:** Department of Ecology and Evolutionary Biology & Biodiversity Institute, University of Kansas, Lawrence, KS 66045 USA; Fish and Wildlife Conservation Department, Virginia Tech, Blacksburg, VA 24060 USA; Research Group of Emerging and Neglected Diseases, Ecoepidemiology and Biodiversity, Health Science Faculty, School of Biomedical Sciences, Universidad Internacional SEK (UISEK), Quito, Ecuador; School of Biological Sciences, Center for Earth Observation and Modeling, University of Oklahoma, Norman, OK, United States of America; Biology Department, Pittsburg State University, Pittsburg, KS 66762 United States of America; Department of Veterinary Pathobiology, College of Veterinary Medicine, Oklahoma State University, Stillwater, Oklahoma, USA

## Abstract

In this study, we explore a suite of new ecological niche modeling approaches to illuminate the distributional potential and distributional dynamics of questing individuals of four tick species in the central Great Plains region. Specifically, we improve on typical approaches in distributional ecology by (1) assigning time-specific environmental values to sampling events so that repeated sampling at the same points yields additional information, (2) using both positive and negative records of the species as inputs in model development, and (3) constraining model response shapes in models to resemble the shapes of fundamental ecological niches. Model outcomes demonstrate both seasonal dynamics and differences between species in terms of the geographic potential of questing behavior by Great Plains tick species. These improved approaches in distributional ecology have considerable potential to enrich distributional understanding of vector species distributions.

## Introduction

Across North America, ticks pose significant threats to the health of humans and animals as vectors of disease agents (Eisen et al. 2017). Tick surveillance programs have demonstrated expansions of tick species’ geographic distributions, and have increased awareness of the ubiquity of tick-borne pathogens over the last few decades (Sonenshine 2018). In some areas, such as in the central Great Plains—defined as Kansas and Oklahoma for the purposes of this paper—tick diversity and abundance vary geographically between east and west or north and south; these contrasts, at least in part, are consequences of tick host preferences and changing tick habitats (Wimms et al. 2023, Ng’eno et al. 2024). Peripheral regions within tick species’ ranges, like the central Great Plains, offer opportunities to understand tick range and activity dynamics as expansion occurs across the region.

In the central Great Plains, Oklahoma and Kansas specifically, several established and medically-relevant tick species are present, including *Amblyomma americanum, A. maculatum, Dermacentor albipictus, D. variabilis*, and *Ixodes scapularis* (Mitcham et al. 2017). These species differ in their distribution, host preferences, peak activity periods, habitats, and the pathogens that they vector (Saleh et al. 2021); all of these variables must be considered when assessing overall tick risk. For instance, adult *A. americanum* and *I. scapularis* prefer white-tailed deer (*Odocoileus virginianus*) as hosts and forests as habitat, but their peak activity periods are distinct, in spring-summer or fall-winter, respectively (Schulze et al. 2001, Noden et al. 2022). Therefore, risk for *I. scapularis*-vectored pathogens will be higher in fall and winter, whereas *A. americanum*-vectored pathogens will be of greater concern in spring and summer.

In this contribution, we develop models to produce detailed “time-and-space” characterizations of environmental conditions that favor questing activity for four medically important tick species that are found across Kansas and Oklahoma (*A. maculatum, D. albipictus, D. variabilis*, and *I. scapularis*). We have previously presented models for *A. americanum*, the most common (i.e., massively dominant) species in tick communities in the region (Cobos et al. 2024, Ng’eno et al. 2024). This suite of models is novel, assessing heterogeneity in distributions of questing ticks in both time and space, using time-specific environmental signatures and explicitly considering sampling effort. Here, we analyze the remaining four tick species that we encountered in our two-year (2020-2022) field program, including 181 visits to 10 sites across central and eastern Kansas and Oklahoma (Ng’eno et al. 2024). This analysis illustrates two important points: (i) the impressive value of time-specific analyses in increasing sample sizes when sampling is conducted repeatedly at relatively few sites, and (ii) the diversity of phenology and activity patterns among different tick species in the central Great Plains region. This contribution constitutes a first step toward a synthetic view of risk of transmission of tick-borne pathogens to humans, both across space and through time across a broad region.

## Methods

The analyses presented herein were developed in four main steps, as follows. (i) Prepare occurrence and sampling records for each tick species using information obtained from detailed field sampling during 2020–2022; (ii) develop weekly environmental data summaries to include time-specific information on conditions across the study area while associating those conditions with tick occurrence and sampling records; (iii) test for sets of environmental conditions that are related to occurrence and non-occurrence of each tick species; and finally, (iv) develop ecological niche models and use them to create weekly predictions for each tick species. Scripts to reproduce all analyses are provided at https://github.com/marlonecobos/Tick_KSOK/tree/main/Four_species_ENM.

### Presence-absence data

Occurrence (presence) and non-occurrence (apparent absence) data were obtained from a program of longitudinal sampling at 10 sites across Kansas and Oklahoma, lasting between 1 August 2020 and 30 September 2022 (Fig 1), and described in detail elsewhere (Ng’eno et al. 2024). The sampling included intensive efforts to collect questing ticks in wooded and open areas at each site in the study, using a consistent sampling protocol combining flagging, dragging, and CO_2_ traps. Sampling teams to each site consisted of 3–7 persons, and sampling visits lasted 3–4 hours. Sampling was conducted under conditions considered to be appropriate for tick questing (temperature 2–32°C, wind speed <15 mi/h, and no precipitation or dew). In all, sampling effort comprised 181 visits across the 10 sites.

**Fig 1.**
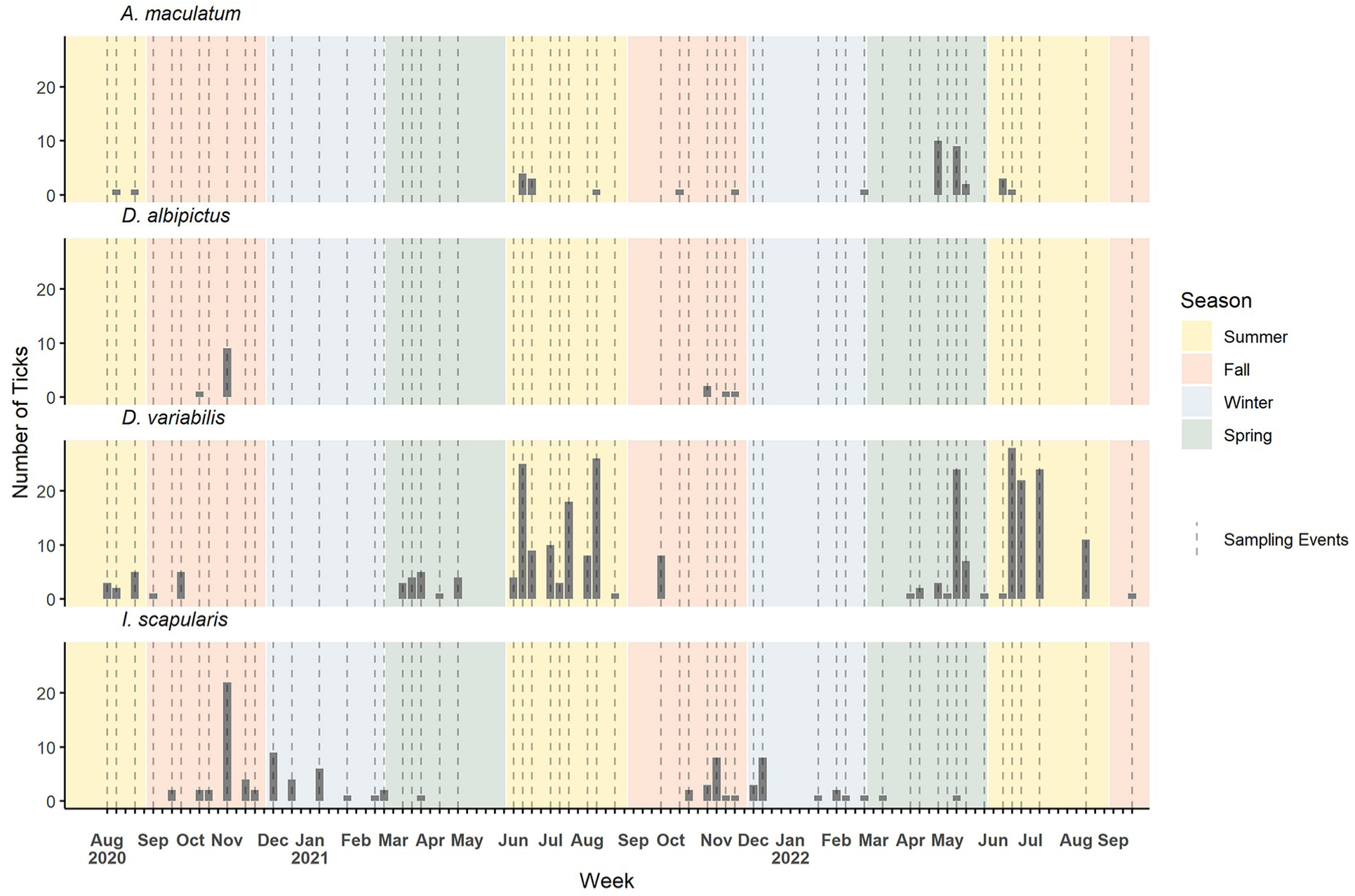
Summary of sampling events (vertical dashed lines) and detections of ticks of four species (*Amblyomma maculatum*, *Dermacentor albipictus*, *D. variabilis*, and *Ixodes scapularis*; gray bars) across the 10 sites sampled as part of our field studies of questing tick behavior. Seasons are shown as green = Spring, yellow = Summer, red = Fall, and blue = Winter.

From the set of days and sites sampled (Fig 1), we obtained records of presences and absences for each of the four tick species to be treated in this paper (models for *A. americanum* have been presented separately; Cobos et al. 2024). For *D. variabilis*, we collected 289 (78.1%) adults and 42 (11.4%) nymphs, so models were developed only for adults of this species. Only adult ticks were collected for *A. maculatum* and *I. scapularis*; only larvae were collected for *D. albipictus*. We assembled data sets in the form of all records of each of the species, to allow analysis of the four species. A presence record is a unique combination of tick species, day, month, year, and site; an absence is also a unique combination of the same fields, but for which the species was not detected. All data processing steps were performed using base functions in R (R Core Team 2023).

### Environmental data

We obtained climatic data from the Daymet database (Daymet 2024). We included the following variables in analyses: maximum air temperature (tmax), minimum air temperature (tmin), precipitation (prcp), shortwave radiation (srad), water vapor pressure (wvap), and day length (dayl). The snow water equivalent variable, also available in these datasets, was excluded because none of our field sampling occurred under conditions with snow cover; consequently, any analyses of conditions with nonzero values of this variable would be extrapolative, and indeed this variable caused considerable confusion in results when included in preliminary exploratory analyses.

The summary climatic layers at approximately 1 km spatial resolution were cropped to a rectangular extent (33.62–40.00°N, 103.00–94.5°W) covering entire the states of Kansas and Oklahoma. We produced environmental data layers at a temporal resolution of 8 days, averaging over daily values, for the entire span of 1980–2022, although we analyze only the years 2020-2022 in this contribution, as they correspond temporally to the sampling events that produced the tick presence-absence data. The 8-day averaging periods resulted in 46 such sets of data layers per year (i.e., Julian day 1-8, 9-16, …), although the last layer of each year covered only 5-6 days, depending on the year; we used the Julian day for each 8-day average layer for naming layers uniquely. Variable processing analyses were done in Google Earth Engine (Gorelick et al. 2017).

In a basic exploration of relative independence versus dependence among environmental variables, we calculated Pearson’s correlation coefficients among the 6 original variables, and detected high correlations (i.e., | *r* | > 0.8) between several pairs of variables (S1 Table). As a consequence, we used principal components analysis (PCA) to obtain sets of orthogonal predictors from the original variables (S2 and S3 Tables). This analysis was applied to data covering January 2020 through December 2022, and we used the PCA to characterize every Julian week (8-day period) over the three years of analysis. We used the tick presence and absence occurrence record sets described above to extract values from the principal components according to the geographic coordinates and the Julian week of collection. This step created a dataset that characterized occurrences and non-occurrences in our tick-collection data in environmental dimensions specific in both time and space. All of these raster processing procedures were performed with the R package terra (Hijmans 2022).

### Niche signal exploration

A first analysis step was to test for a niche signal—in effect, a non-random distribution in environmental space as regards which of our sampling events did *versus* did not detect the tick species in question. If the species is present in all samples, or if detections are random with respect to environmental dimensions, then no niche signal is manifested across that time span and study region. Cobos and Peterson (2022) presented multivariate (PERMANOVA) and randomization-based univariate tests for such niche signals. The PERMANOVA tests assess whether the position and dispersion of the records of detection are similar to all of the records (detections and non-detections) (Cobos and Peterson 2022). Rejecting the PERMANOVA null hypothesis indicates that detections show a different environmental signal compared to all records (the sampling universe).

The univariate tests, on the other hand, assess niche signals via random resampling of the overall dataset to assess the random-distribution null hypothesis directly (Cobos and Peterson 2022). Specifically, if one has an overall sample of size *N*, out of which a smaller subset of samples (termed *x*) proved to be positive for a phenomenon (in this case, questing behavior by a particular tick species), then a large number (e.g., in this case 1000) of samples of size *x* are drawn at random from the full set of *N* sampling events to create a null distribution. The mean, median, standard deviation, and range of these replicate samples in terms of environmental data values are then characterized, and the observed value among the real *x* positive samples is compared to that distribution. Niche signal explorations were performed using the R package enmpa (Cobos and Peterson 2022)(Arias-Giraldo and Cobos 2024).

### Ecological niche model development

Ecological niche model calibration was done via methods that contrast rather markedly with the usual approach to such questions (see, e.g., Peterson et al. 2011), in view of the availability of high-quality presence and absence data from our tick sampling program across the two states (detection and non-detection of questing ticks). We used logistic generalized linear models (GLMs), which contrast populations of positive (detections) and negative (non-detections) occurrence data in predictions of probability of “presence” *versus* “absence.” GLMs based on contrasts between presence data and pseudoabsence data resampled from the parts of the study area that did not hold presences are not appropriate in ecological niche modeling (Peterson et al. 2011); however, they can be appropriate and powerful approaches when high-quality presence-absence data are available (Cobos et al. 2024).

Our GLM calibration approach consisted of testing a large suite of candidate models representing all possible combinations of predictors derived from linear and quadratic responses of 6 principal component variables, for a total of 4095 candidate models. All candidate models were created and evaluated for predictive ability using a 10 *k*-fold partitioning approach (i.e., the method to split training and testing data) (Peterson et al. 2011). Two metrics, area under the curve of the receiver operating characteristic (ROC AUC; Elith et al. 2006) and true skill statistic (TSS; Allouche et al. 2006) were used to assess performance and predictive power of candidate models; note that use of these metrics is appropriate only in cases in which positive and negative data are available (Lobo et al. 2008). We also created candidate models with the whole set of data and used the Akaike information criterion (AIC; Warren and Seifert 2011) to assess model goodness-of-fit penalized by model complexity.

Once all candidate models were fitted and evaluated, we retained the subset of models that passed four filtering steps, following pipelines presented in previous approaches to presence-only niche models (Cobos et al. 2019). That is, we retained models (1) that had all quadratic predictor coefficients negative (i.e., variable response curves were convex or unimodal, and not concave or bimodal); from among those models, we retained (2) models that had AUC > 0.5, and then kept only (3) those with TSS > 0.4. Finally, (4) we chose as final models only the ones with AIC scores within 2 units of the minimum value among those that had passed the first three filters (i.e., ΔAIC ≤ 2).

Once we had selected final models, we transferred each of the set of best models separately to all weekly environmental summaries for the sampling period. We created a weighted average of all best model predictions to obtain a single consensus prediction per week based on the AIC weights calculated for selected models. All modeling steps were performed using the R package enmpa (Arias-Giraldo and Cobos 2024).

### Post-modeling analyses

Given the vagaries of weather and its variation in individual days and weeks, we smoothed weekly predictions using a moving window approach, in which each week was replaced with the average value of the week before, the focal week, and the following week (Cobos et al. 2024). We also produced monthly averages from weekly predictions by creating averages of the weekly rasters across the months in which they fall, including partial membership of weeks that overlap between consecutive months.

To identify areas in Kansas and Oklahoma with environmental conditions outside the ranges of conditions represented in the data used for models, we used the mobility-oriented parity metric (MOP; Owens et al. 2013). MOP analyses were applied to all environmental summaries for 2020–2022. Monthly summaries of MOP results were obtained similarly to monthly averages of model predictions; higher values in these summaries indicate that such areas have been outside calibration ranges for more of the month. MOP analyses were done using the R package mop (Cobos et al. 2023). All other analyses in this section were done using the R package terra.

## Results

### Niche signal tests

The univariate tests for non-random distribution of detections of each species with respect to environmental conditions indicated significant differences from null expectations in most (but not all) of the environmental dimensions that we assessed (Fig 2). Some were as expected, given the phenological variation among species that we have documented elsewhere (Ng’eno et al. 2024): for example, the two fall-winter tick species (i.e., *I. scapularis*, *D. albipictus*) are found questing under conditions of shorter day length, whereas the two spring-summer tick species (i.e., *D. variabilis*, *A. maculatum*) are found questing under longer day length, as compared with null expectations from the sampling scheme.

**Fig 2.**
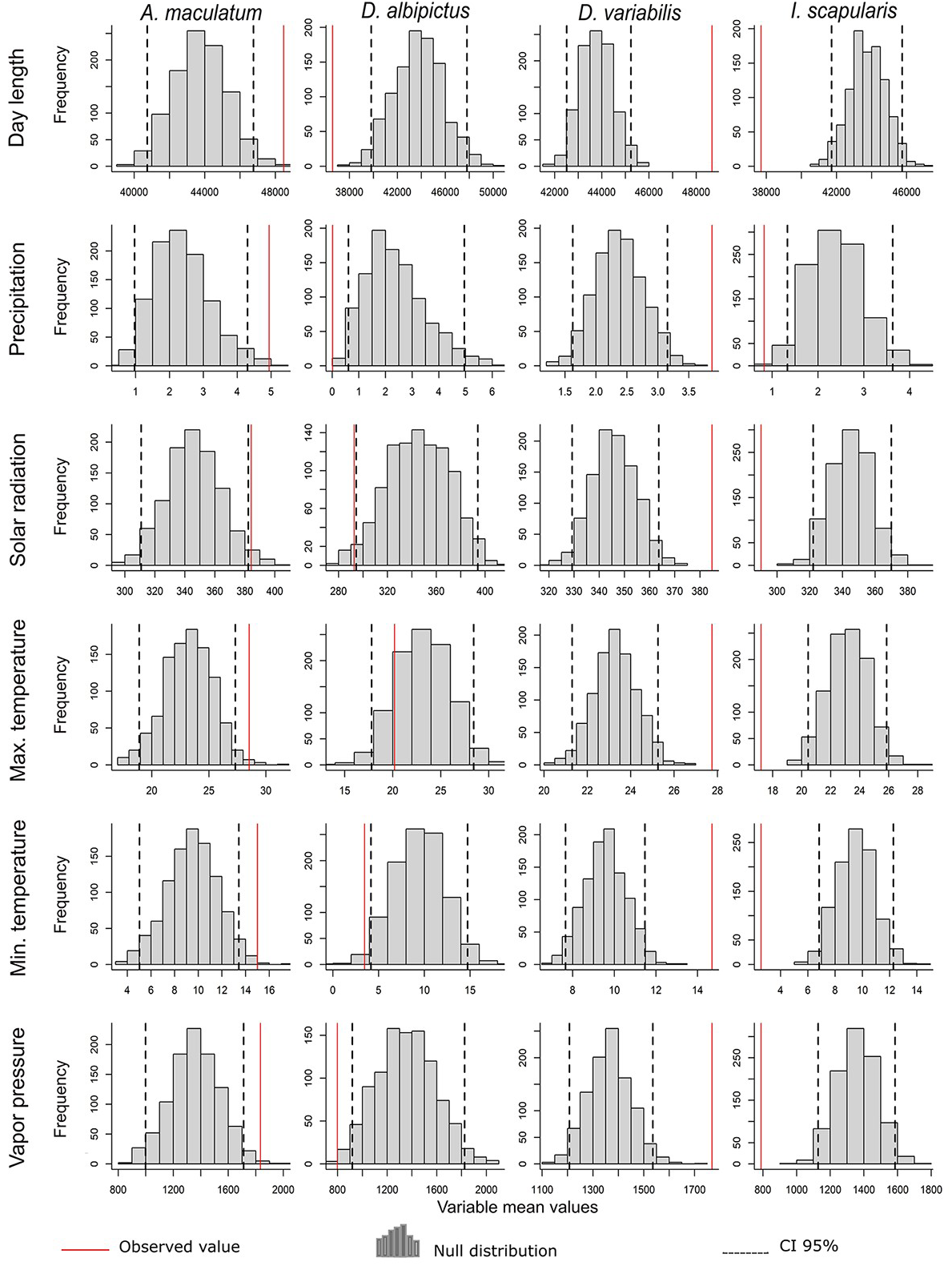
Summary of univariate tests of niche signal (i.e., non-random distribution with respect to availability of sampled conditions) for four tick species with respect to six climatic parameters. Gray bars indicate frequency of values among 1000 null resamplings from the available data (see Methods); black vertical dashed bars indicate the 2.5th and 97.5th percentiles of the null distribution. Vertical red line indicates the observed value.

In fact, these same differences hold for precipitation and vapor pressure, with fall-winter ticks occurring under drier conditions and spring-summer ticks occurring under wetter conditions compared to null expectations. The relationships of the two temperature variables and solar radiation showed similar contrasts between fall-winter and spring-summer tick species, but the rarer species (*D. albipictus*, *A. maculatum*) were more equivocal in the significance of their niche signal, likely owing to lower statistical power.

### Ecological niche models

The four species under analysis in this contribution show at least two distinct signatures of seasonality of questing behavior, as can be appreciated in the form of Spring-Summer peaks of suitability for *A. maculatum* and *D. variabilis*, contrasting with Fall peaks in *D. albipictus* and Fall-Winter peaks in *I. scapularis* (analyzed in detail elsewhere; Ng’eno et al. 2024). These seasonal patterns of variation in suitability, as reflected in model suitability signatures through time, echo the phenology of the species (Fig 4; Ng’eno et al. 2024). Of particular note is the covariation between suitability patterns and key climatic variables, as is illustrated in Fig 3 as well. These seasonal patterns form the basis for the model-based predictions that are the centerpiece of this contribution.

**Fig 3.**
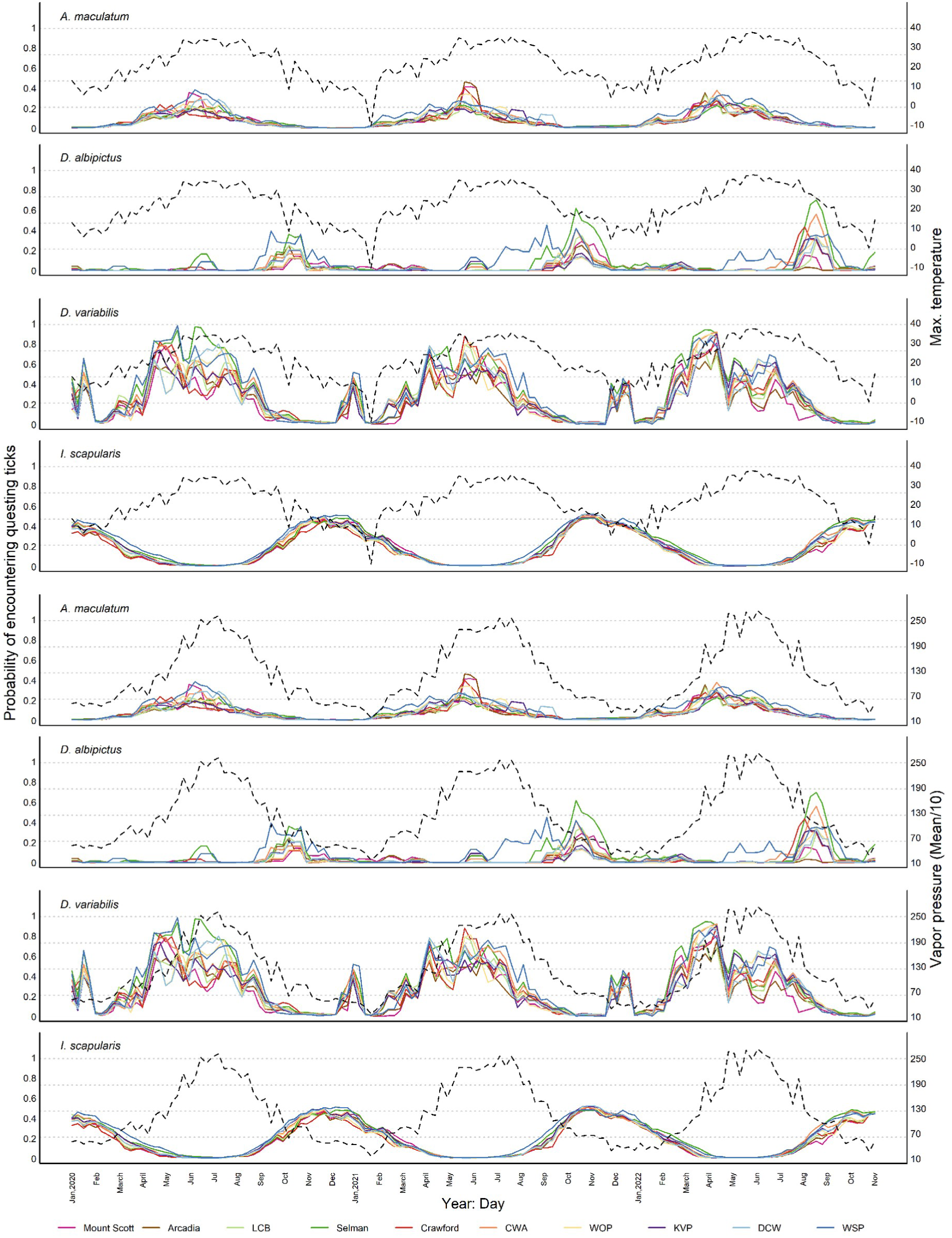
Temporal traces across all of 2020-2022 for suitability for four species of ticks at 10 sites in Kansas and Oklahoma. Suitability at each site is shown as a solid line with a different color; average maximum temperature and vapor pressure across all sites are shown as a dotted line for illustration of environmental trends in the region.

**Fig 4.**
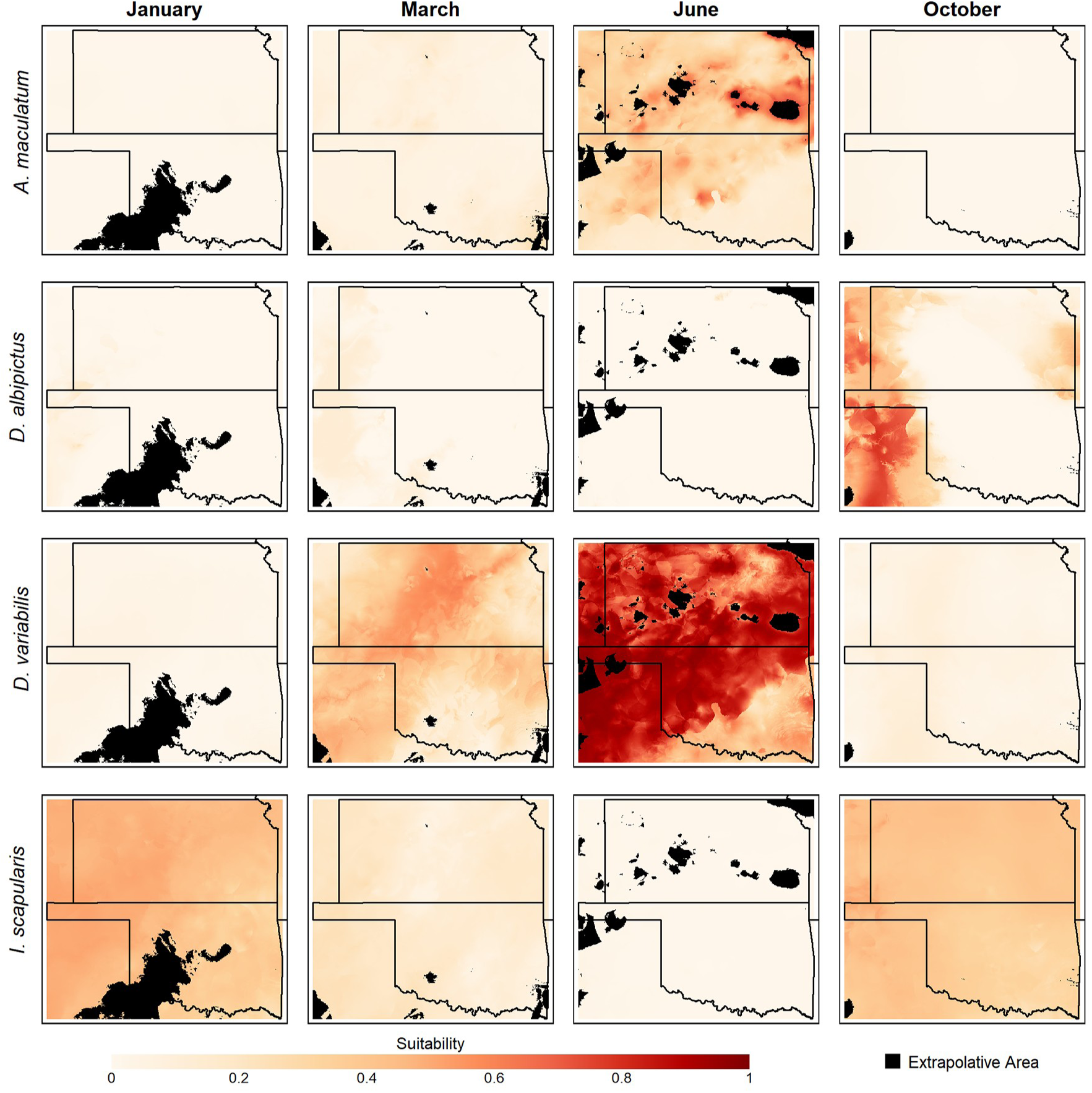
Summary of time-specific ecological niche model predictions for four example weeks in 2021, for four tick species. Areas shown in black were identified as presenting extrapolative conditions in that time period via MOP analyses. The remaining areas were classified according to modeled suitability, ranging from nil (white) to highly suitable (dark red).

The ecological niche models developed for the four tick species reflected these same contrasts between phenological types of tick species (i.e., fall-winter *versus* spring-summer; Fig 4). That is, *D. variabilis* begins to rise in activity in Spring, and reaches a maximum in the Summer; *A. maculatum* is more strictly focused in Summer. Similarly, larvae of *D. albipictus* are active in the Fall, whereas *I. scapularis* is active in Fall and Winter. Extrapolative conditions were found in southwestern Oklahoma in Winter in terms of PC6, and were in scattered parts of Kansas and western Oklahoma in Summer in terms of different combinations of PC1, PC2, PC3, PC5, and PC6 (Fig 4; also see supplemental materials). Overall, areas presenting extrapolative conditions were generally small, and did not cover much of the study area.

## Discussion

### New modeling approaches

This study explores the potential strengths of a suite of new ecological niche modeling approaches, as applied to four tick species in the central Great Plains region. We incorporate three major changes from the typical analytical approaches in distributional ecology: (1) time-specific environmental values assigned to individual sampling events, (2) use of information directly from the sampling that produced the positive and negative records of the species, and (3) constraint of the response shapes in the models to be convex and more consistent with current understanding of what fundamental ecological niches should be like. More detail is provided below on each of these points.

Typically, ecological niche modeling applications use a single environmental value for each site, neglecting the sometimes-extreme environmental variation that is manifested at a site in the course of days, years, or decades (Ingenloff and Peterson 2020). Early explorations of the idea of time-specific niche modeling approaches showed the greater specificity that can be achieved by avoiding the fallacy that an average can represent the variation that goes into the average value adequately (Peterson et al. 2005). A later paper explored contrasts between time-specific and time-averaged approaches in distributional ecology, and provided methodological protocols (although they were computationally intensive) for handling presence-only data in such analyses (Ingenloff and Peterson 2020).

The analyses presented herein improve on those previous time-specific methods by direct and full incorporation of information from the sampling that produced the set of positive records of each species (Cobos et al. 2024). That is, in the analyses presented herein, our models represent statistical contrasts between sampling events that did and did not detect a given species. In this sense, the negative data in our analyses are not “pseudo-absence” data that are assumed to represent absence of the species (Peterson et al. 2011); rather, they are sampling events with field methods equivalent to those that yielded presence records, but on which the species in question was not encountered. In this sense, we have refined the negative data considerably from the usual approach in distributional ecological analyses, which is likely to illuminate the niche signal recovered in the modeling process considerably.

Finally, we pursue yet another improvement to the approach of testing models calibrated under different parameter settings that began with the work of Warren and Seifert (2011), and refined and extended considerably with the development of the KUENM R package (Cobos et al. 2019). That is, for a number of years, model selection has included consideration of model significance and predictive performance, as well as model simplicity. Here, following other recent analyses (e.g., Drake 2015), we add one more consideration: that model response types “look like” fundamental ecological niches, in that they are unimodal with respect to each environmental dimension, as well as convex in multivariate space (Jiménez et al. 2019, Soberón and Peterson 2019, Jiménez and Soberón 2022).

### Diversity of tick questing activity patterns

As witnessed in the present paper, tick questing behavior varies across time, space, and by species. Ticks are known to prefer particular climatic variables (e.g., relative humidity, ambient temperature), but can survive weather extremes (Nicholson et al. 2019). The latter scenario is critical for long-term tick survival; however, their preferences dictate their peak activity or questing periods, which relate or correspond to times at which interaction of ticks with hosts may be highest, which in turn drives spatiotemporal patterns of tick-borne disease risk. Across the four species presented here, variation in questing patterns was dramatic, which was unsurprising based on prior research in other regions of North America and the seasonal variation expected for the central Great Plains (Raghavan et al. 2007, Saleh et al. 2019, Cull 2022).

For warm-season ticks, such as *D. variabilis* and *A. maculatum*, their questing periods peaked in the summer months, in general. These two species are known to enter diapause in cooler months of the year, overwintering as immature stages; they may have an extended life cycle duration in comparison to other tick species. Once temperatures rise above ∼20°C, immature stages of these species are more likely to be questing for hosts; temperature known to be an important predictor of their survival (Fieler et al. 2021). In contrast, cooler-season ticks (*I. scapularis* and larval *D. albipictus*), have peak activity periods in late fall or winter, although even these ticks will pause activity under conditions of more extreme weather (Holmes et al. 2018).

In the central Great Plains, these temperature-dependent behaviors are expected to vary based on specific location. Prior work has demonstrated that environmental conditions in some parts of this region support ticks year-round (Sundstrom et al. 2021), which is not the case across the entirety of the central Great Plains (Noden et al. 2022, CDC 2024). As certain vegetation types (e.g., forest) and reservoir hosts (e.g., deer) are increasingly able to survive in historically non-endemic tick regions of the central Great Plains, novel tick populations may enter and establish (Noden et al. 2021, Noden et al. 2024), or perhaps they have been there, but at low population levels, and are now able to increase. Most of the study sites in the present study were in the eastern or central regions of the two states; however, if the westernmost parts of Oklahoma and Kansas are considered, temperature, relative humidity, and terrain are quite different. More work is warranted to compare and identify additional factors that can lead to tick expansion in this region so more accurate risk modeling can occur for areas with emerging populations of medically-relevant ticks.

### Conclusion and public health implications

This paper presents a complex and varied view of tick questing activity through the year and across the central Great Plains, across four tick species. The fifth tick species that is widespread in the region, *A. americanum* is treated in a separate contribution (Cobos et al. 2024). This complexity of questing activity by a diversity of tick species has important implications for public health in the region. A large subsample of the same ticks that were collected as part of the work reported herein have been tested for a battery of bacterial pathogens (e.g., *Borrelia*, *Anaplasma*, *Ehrlichia*); results of those tests with respect to pathogens will be reported elsewhere (K. Duncan et al. in prep.). Overall, the picture is one of extreme complexity of multiple species of pathogens transmitted by multiple species of ticks, which in turn have diverse life histories, ecological niches, phenologies, seasonalities, and spatiotemporal distributional patterns.

## Acknowledgments

This study was made possible thanks to the hard work of the Central Great Plains Tick-borne Disease Risk team, and the analyses were greatly facilitated by work and advice from the KUENM Working Group (University of Kansas).

## Data accessibility statement

Tick data analyzed in this project and the code to reproduce all analyses are openly available at https://github.com/marlonecobos/Tick_KSOK. Other data used can be accessed from the sources described in the Methods section.

## Supplementary Information

**Table S1.** shows the Pearson product moment correlations among environmental variables. Correlations greater than 0.5 or less than -0.5 are indicated in bold.

|  | Mean day length | Mean precipitation | Mean solar radiation | Mean maximum temperature | Mean minimum temperature | Mean vapor pressure |
| --- | --- | --- | --- | --- | --- | --- |
| Mean day length | 1 | – | – | – | – | – |
| Mean precipitation | 0.28 | 1 | – | – | – | – |
| Mean solar radiation | **0.75** | -0.13 | 1 | – | – | – |
| Mean maximum temperature | **0.73** | -0.001 | **0.73** | 1 | – | – |
| Mean minimum temperature | **0.82** | 0.15 | **0.64** | **0.95** | 1 | – |
| Mean vapor pressure | **0.80** | 0.12 | **0.58** | **0.91** | **0.97** | 1 |

**Table S2.** shows the raw climatic variable loadings on each of the 6 principal components.

|  | PC1 | PC2 | PC3 | PC4 | PC5 | PC6 |
| --- | --- | --- | --- | --- | --- | --- |
| Mean day length | 0.456 | -0.026 | 0.379 | 0.634 | 0.448 | -0.211 |
| Mean precipitation | 0.121 | 0.836 | 0.461 | -0.235 | -0.139 | -0.014 |
| Mean solar radiation | 0.359 | -0.492 | 0.553 | -0.194 | -0.502 | 0.183 |
| Mean maximum temperature | 0.474 | -0.099 | -0.189 | -0.597 | 0.255 | -0.556 |
| Mean minimum temperature | 0.482 | 0.098 | -0.262 | -0.138 | 0.293 | 0.765 |
| Mean vapor pressure | 0.438 | 0.199 | -0.483 | 0.361 | -0.614 | -0.167 |

**Table S3.** shows the summary of the variance explained by each of the principal components.

|  | PC1 | PC2 | PC3 | PC4 | PC5 | PC6 |
| --- | --- | --- | --- | --- | --- | --- |
| Standard deviation | 2.004 | 1.076 | 0.762 | 0.402 | 0.264 | 0.120 |
| Proportion of Variance | 0.670 | 0.193 | 0.097 | 0.027 | 0.012 | 0.002 |
| Cumulative Proportion | 0.670 | 0.862 | 0.959 | 0.986 | 0.997 | 1 |

**Tables S4-S7.** show the summaries of the univariate tests for the existence of a distinguishable environmental bias (i.e., ecological niche) of the four focal tick species. “Higher” and “lower” indicate that the observed value of the tick occurrences with regard to the relevant environmental variable is located in the upper or lower 2.5% of the null distribution, respectively. **S4**. *Amblyomma maculatum*

| Variable | H0 mean | H0 median | H0 SD | H0 range | Positive mean vs null | Positive median vs null | Positive SD vs null | Positive range vs null |
| --- | --- | --- | --- | --- | --- | --- | --- | --- |
| Mean day length | rejected | rejected | accepted | accepted | higher | higher | indistinct | indistinct |
| Mean precipitation | rejected | rejected | accepted | rejected | higher | higher | indistinct | higher |
| Mean solar radiation | rejected | accepted | accepted | accepted | higher | indistinct | indistinct | indistinct |
| Mean maximum temperature | rejected | rejected | accepted | accepted | higher | higher | indistinct | indistinct |
| Mean minimum temperature | rejected | rejected | accepted | accepted | higher | higher | indistinct | indistinct |
| Mean vapor pressure | rejected | rejected | accepted | accepted | higher | higher | indistinct | indistinct |

**S5.** Dermacentor albipictus.

**S5. *Dermacentor albipictus***
| Variable | H0 mean | H0 median | H0 SD | H0 range | Positive mean vs null | Positive median vs null | Positive SD vs null | Positive range vs null |
| --- | --- | --- | --- | --- | --- | --- | --- | --- |
| Mean day length | rejected | rejected | rejected | rejected | lower | lower | lower | lower |
| Mean precipitation | rejected | accepted | rejected | rejected | lower | indistinct | lower | lower |
| Mean solar radiation | rejected | accepted | rejected | rejected | lower | indistinct | lower | lower |
| Mean maximum temperature | accepted | accepted | rejected | rejected | indistinct | indistinct | lower | lower |
| Mean minimum temperature | rejected | accepted | rejected | rejected | lower | indistinct | lower | lower |
| Mean vapor pressure | rejected | accepted | rejected | rejected | lower | indistinct | lower | lower |

**S6.** Dermacentor variabilis.

**S6. *Dermacentor variabilis***
| Variable | H0 mean | H0 median | H0 SD | H0 range | Positive mean vs null | Positive median vs null | Positive SD vs null | Positive range vs null |
| --- | --- | --- | --- | --- | --- | --- | --- | --- |
| Mean day length | rejected | rejected | rejected | rejected | higher | higher | lower | lower |
| Mean precipitation | rejected | rejected | accepted | accepted | higher | higher | indistinct | indistinct |
| Mean solar radiation | rejected | rejected | rejected | accepted | higher | higher | lower | indistinct |
| Mean maximum temperature | rejected | rejected | rejected | rejected | higher | higher | lower | lower |
| Mean minimum temperature | rejected | rejected | rejected | rejected | higher | higher | lower | lower |
| Mean vapor pressure | rejected | rejected | rejected | rejected | higher | higher | lower | lower |

**S7.** Ixodes scapularis.

**S7. *Ixodes scapularis***
| Variable | H0 mean | H0 median | H0 SD | H0 range | Positive mean vs null | Positive median vs null | Positive SD vs null | Positive range vs null |
| --- | --- | --- | --- | --- | --- | --- | --- | --- |
| Mean day length | rejected | rejected | rejected | rejected | lower | lower | lower | lower |
| Mean precipitation | rejected | accepted | rejected | accepted | lower | indistinct | lower | indistinct |
| Mean solar radiation | rejected | rejected | rejected | accepted | lower | lower | lower | indistinct |
| Mean maximum temperature | rejected | rejected | rejected | rejected | lower | lower | lower | lower |
| Mean minimum temperature | rejected | rejected | rejected | accepted | lower | lower | lower | indistinct |
| Mean vapor pressure | rejected | rejected | rejected | rejected | lower | lower | lower | lower |

**Tables S8-S11.** show “Best” models selected in the process of model calibration and evaluation. “^2” indicates quadratic terms in a model. Values for AUC, sensitivity, specificity, and TSS are averages deriving from the 10-kfold evaluation process. Threshold indicates the value used to test sensitivity and specificity of the model. AUC = area under the receiver operative characteristic curve; TSS = true skill statistic; AIC = Akaike information criterion; ΔAIC = delta AIC. **S8.** *Amblyomma maculatum*

S8. *Amblyomma maculatum*
| ID | Predictors | Threshold<br>Criteria | Threshold<br>SD | Accuracy<br>SD | Sensitivity<br>SD | Specificity<br>SD | Parameters | AIC | $\Delta$ AIC |
| --- | --- | --- | --- | --- | --- | --- | --- | --- | --- |
| 1 | PC1, PC3, PC6 | maxTSS | 0.043 | 0.123 | 0.158 | 0.141 | 3 | 96.109 | 0 |
| 2 | PC1, PC3 | maxTSS | 0.088 | 0.125 | 0.242 | 0.150 | 2 | 96.672 | 0.563 |
| 3 | PC1, PC3, PC4 | maxTSS | 0.092 | 0.198 | 0.211 | 0.231 | 3 | 98.070 | 1.961 |
| 4 | PC1, PC2, PC3, PC6 | maxTSS | 0.079 | 0.145 | 0.158 | 0.166 | 4 | 97.948 | 1.839 |
| 5 | PC1, PC3, PC4, PC6 | maxTSS | 0.062 | 0.143 | 0.158 | 0.162 | 4 | 97.818 | 1.710 |
| 6 | PC1, PC3, PC5, PC6 | maxTSS | 0.083 | 0.137 | 0.211 | 0.157 | 4 | 97.821 | 1.710 |

**S9.**
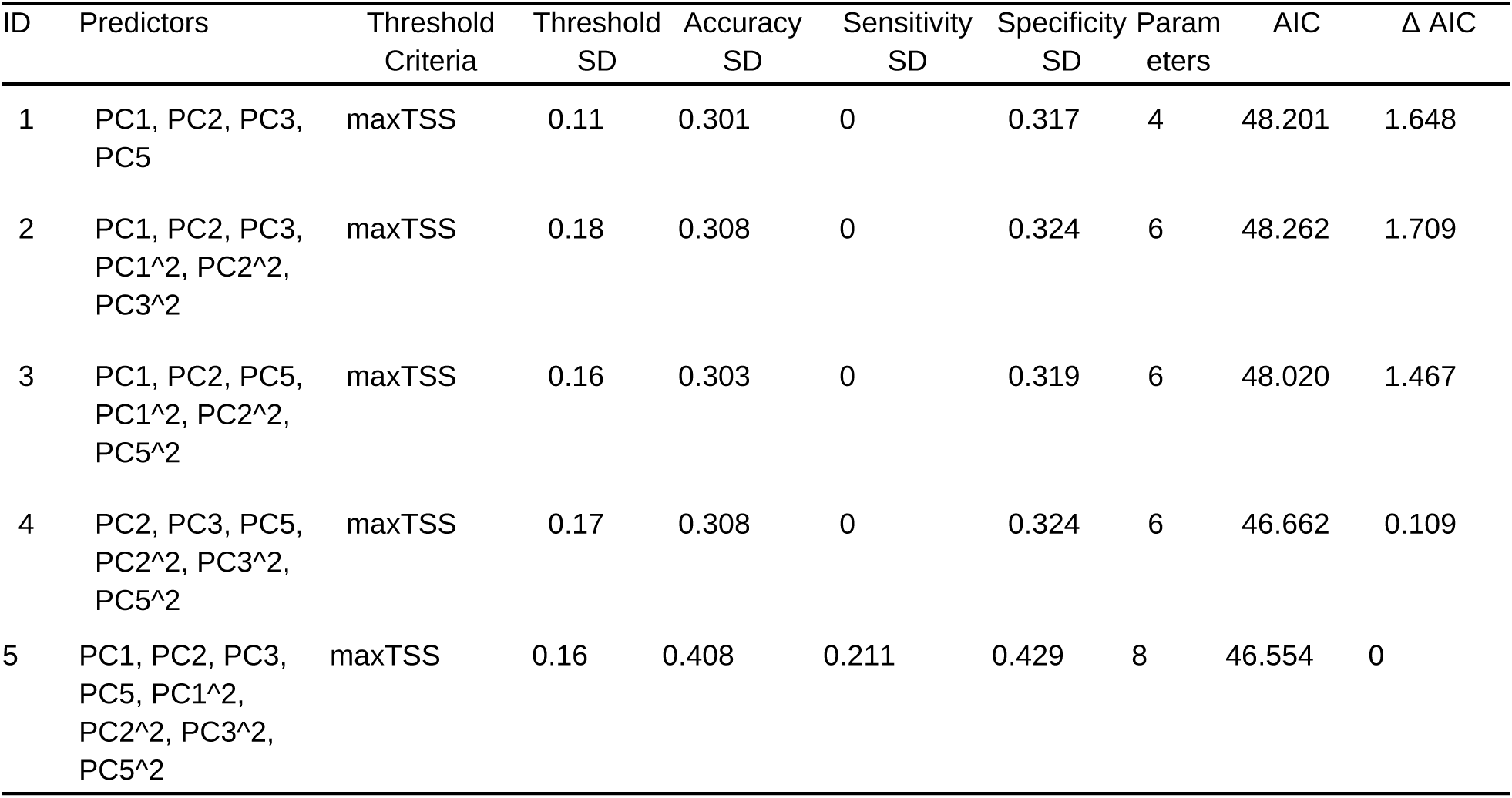
Dermacentor albipictus.

**S10.** Dermacentor variabilis.

**S10. *Dermacentor variabilis***
| ID | Predictors | Threshold | ROC AUC | Sensitivity | Specificity | TSS | AIC | Δ AIC | AIC weight |
| --- | --- | --- | --- | --- | --- | --- | --- | --- | --- |
| 1 | PC1, PC3, PC5, PC4^2, PC6^2 | 0.258 | 0.847 | 0.8833 | 0.776 | 0.659 | 152.78 | 0 | 0.355 |
| 2 | PC1, PC2, PC3, PC5, PC4^2, PC6^2 | 0.227 | 0.847 | 0.9033 | 0.745 | 0.648 | 154.39 | 1.610 | 0.159 |
| 3 | PC1, PC3, PC4, PC5, PC4^2, PC6^2 | 0.183 | 0.828 | 0.9433 | 0.690 | 0.633 | 154.78 | 1.998 | 0.131 |
| 4 | PC1, PC3, PC5, PC6, PC4^2, PC6^2 | 0.218 | 0.843 | 0.920 | 0.721 | 0.641 | 154.16 | 1.381 | 0.178 |
| 5 | PC1, PC3, PC5, PC3^2, PC4^2, PC6^2 | 0.361 | 0.844 | 0.817 | 0.853 | 0.670 | 154.18 | 1.395 | 0.177 |

**S11.** Ixodes scapularis.

**S11. *Ixodes scapularis***
| ID | Predictors | Threshold Criteria | Threshold SD | Accuracy SD | Sensitivity SD | Specificity SD | Parameters | AIC | $\Delta$ AIC |
| --- | --- | --- | --- | --- | --- | --- | --- | --- | --- |
| 1 | PC1, PC3 | maxTSS | 0.1425 | 0.105 | 0.1245 | 0.138 | 2 | 132.8122 | 1.8563 |
| 2 | PC1, PC2, PC3 | maxTSS | 0.0988 | 0.095 | 0.1245 | 0.124 | 3 | 132.6716 | 1.7157 |
| 3 | PC1, PC3, PC4 | maxTSS | 0.1178 | 0.1067 | 0.1245 | 0.137 | 3 | 131.576 | 0.620 |
| 4 | PC1, PC2, PC4 | maxTSS | 0.0874 | 0.1134 | 0.1054 | 0.130 | 3 | 131.161 | 0.205 |
| 5 | PC1, PC3, PC4, PC6 | maxTSS | 0.1367 | 0.1239 | 0.1245 | 0.158 | 4 | 132.915 | 1.959 |
| 6 | PC1, PC2, PC4, PC6 | maxTSS | 0.1408 | 0.1018 | 0.1054 | 0.125 | 4 | 131.971 | 1.016 |
| 7 | PC1, PC2, PC4, PC5 | maxTSS | 0.1306 | 0.1346 | 0.1245 | 0.168 | 4 | 131.889 | 0.933 |
| 8 | PC1, PC2, PC3, PC4 | maxTSS | 0.1213 | 0.1424 | 0.0791 | 0.180 | 4 | 131.437 | 0.481 |
| 9 | PC1, PC2, PC4, PC5, PC6 | maxTSS | 0.1566 | 0.141 | 0.1245 | 0.178 | 5 | 132.614 | 1.658 |
| 10 | PC1, PC2, PC3, PC4, PC6 | maxTSS | 0.1477 | 0.1187 | 0.1245 | 0.156 | 5 | 132.516 | 1.560 |
| 11 | PC1, PC2, PC1<sup>2</sup>, PC2<sup>2</sup> | maxTSS | 0.1233 | 0.0729 | 0.1493 | 0.092 | 4 | 130.956 | 0 |
| 12 | PC1, PC2, PC4, PC1<sup>2</sup>, PC2<sup>2</sup>, PC4<sup>2</sup> | maxTSS | 0.1274 | 0.118 | 0.1245 | 0.145 | 6 | 132.052 | 1.10 |
| 13 | PC1, PC2, PC6, PC1<sup>2</sup>, PC2<sup>2</sup>, PC6<sup>2</sup> | maxTSS | 0.1092 | 0.0946 | 0.1493 | 0.127 | 6 | 132.164 | 1.21 |
| 14 | PC1, PC2, PC4, PC6, PC1<sup>2</sup>, PC2<sup>2</sup>, PC4<sup>2</sup>, PC6<sup>2</sup> | maxTSS | 0.1322 | 0.1121 | 0.1493 | 0.140 | 8 | 132.893 | 1.937 |

**Figures S1-S4.**
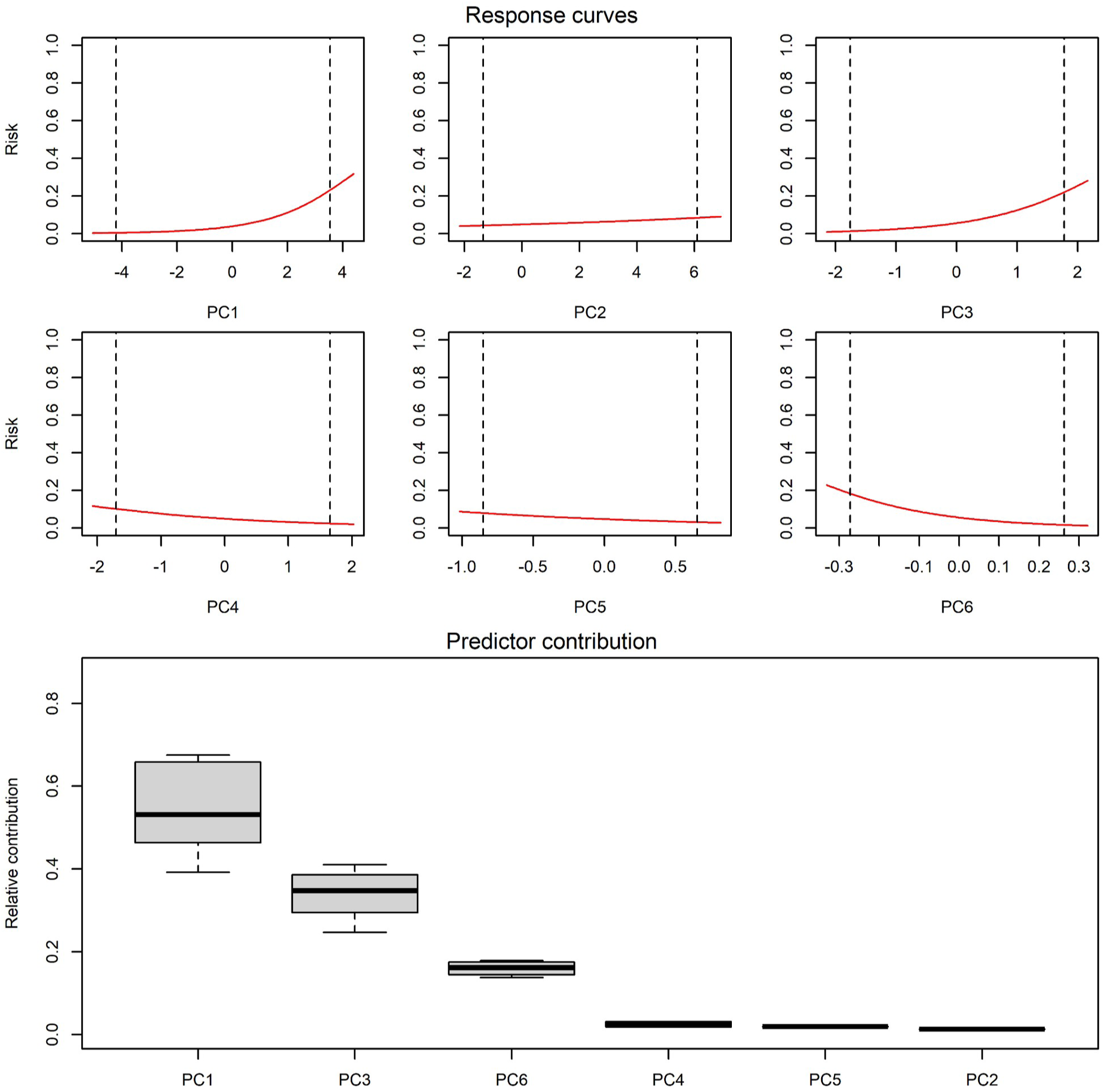
Variable response curves and summary of all forms of predictor relative contribution used in models **S1.** *Amblyomma maculatum*

**S2.**
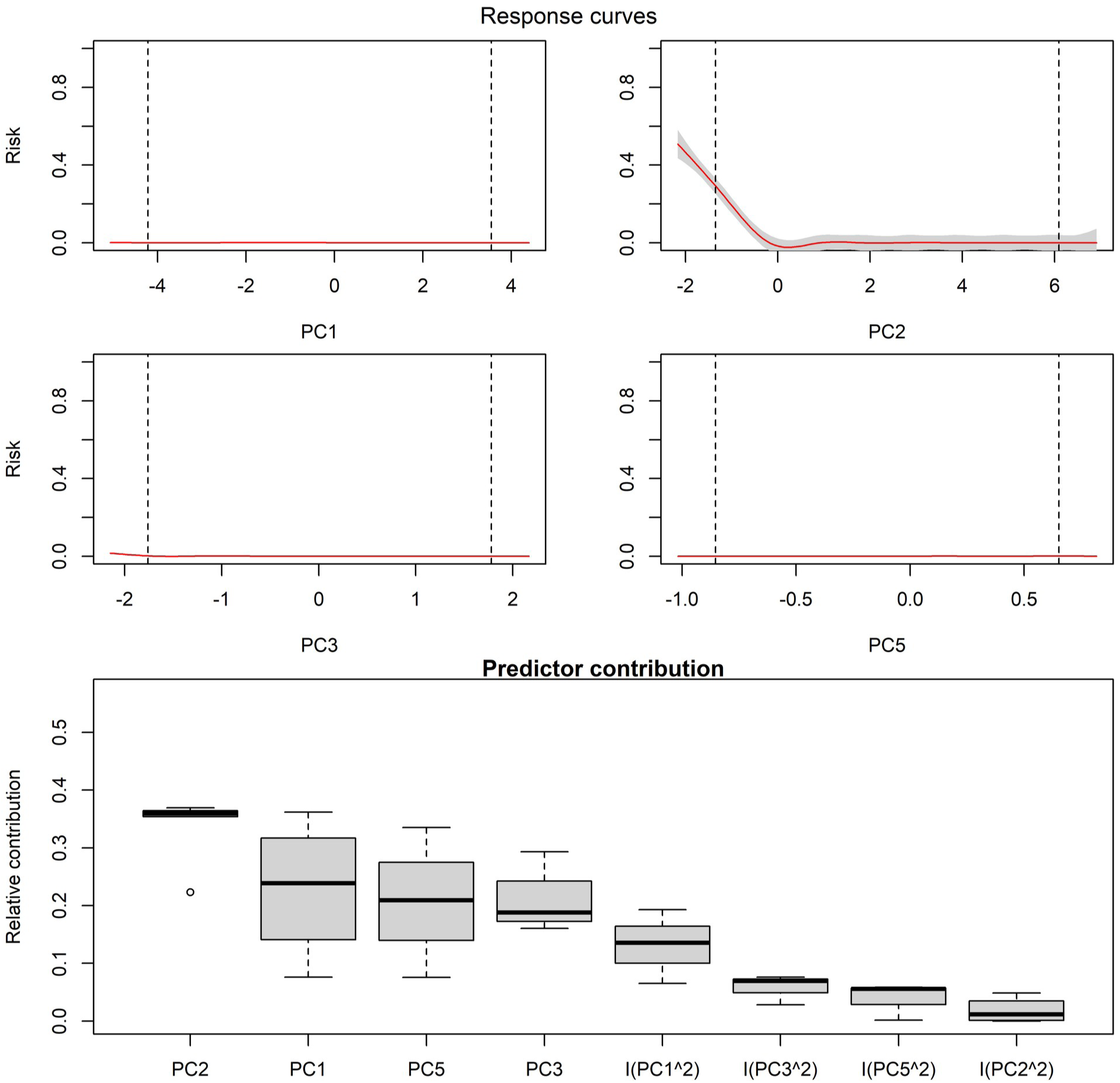
Dermacentor albipictus

**S3.**
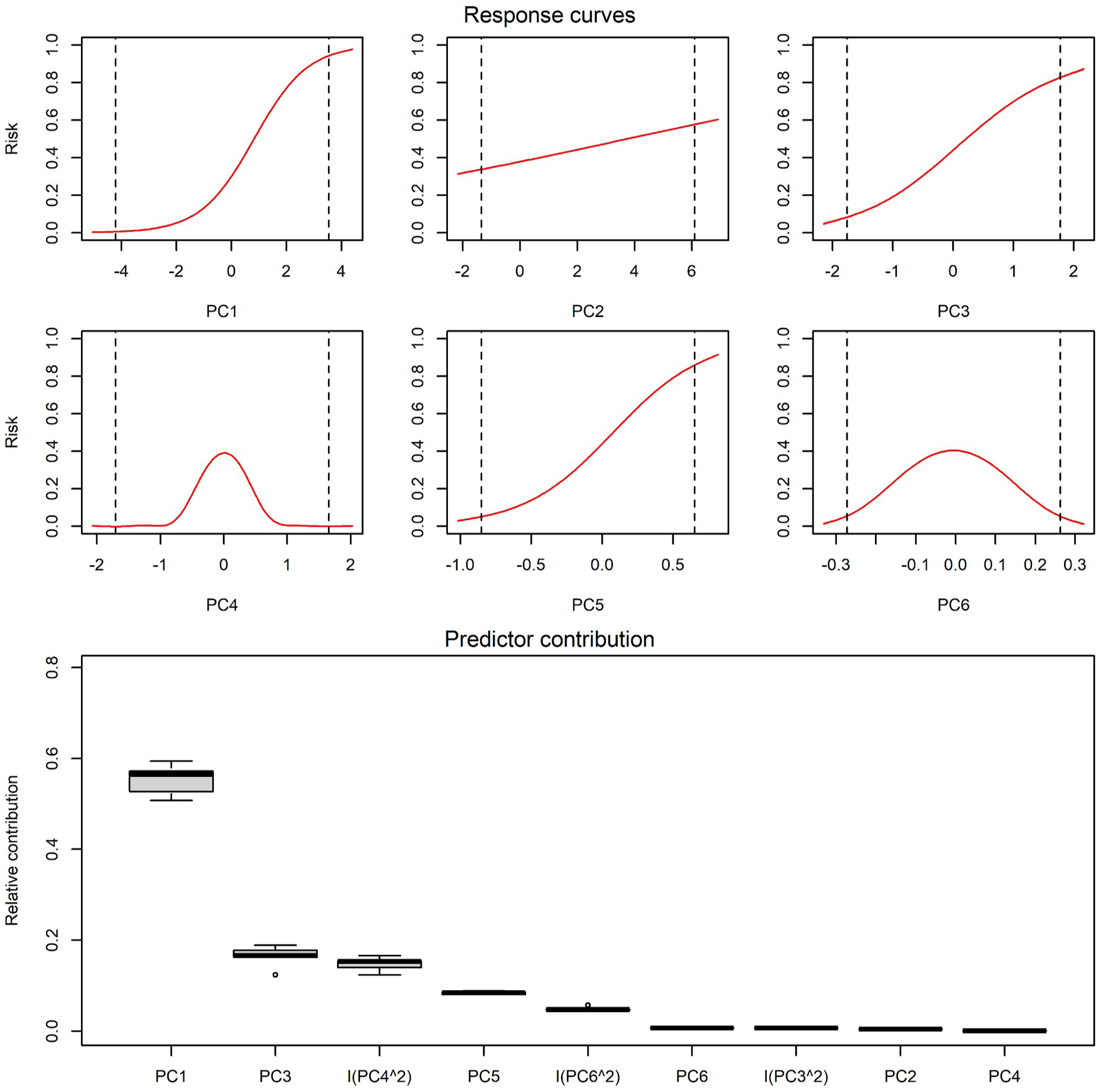
Dermacentor variabilis

**S4.**
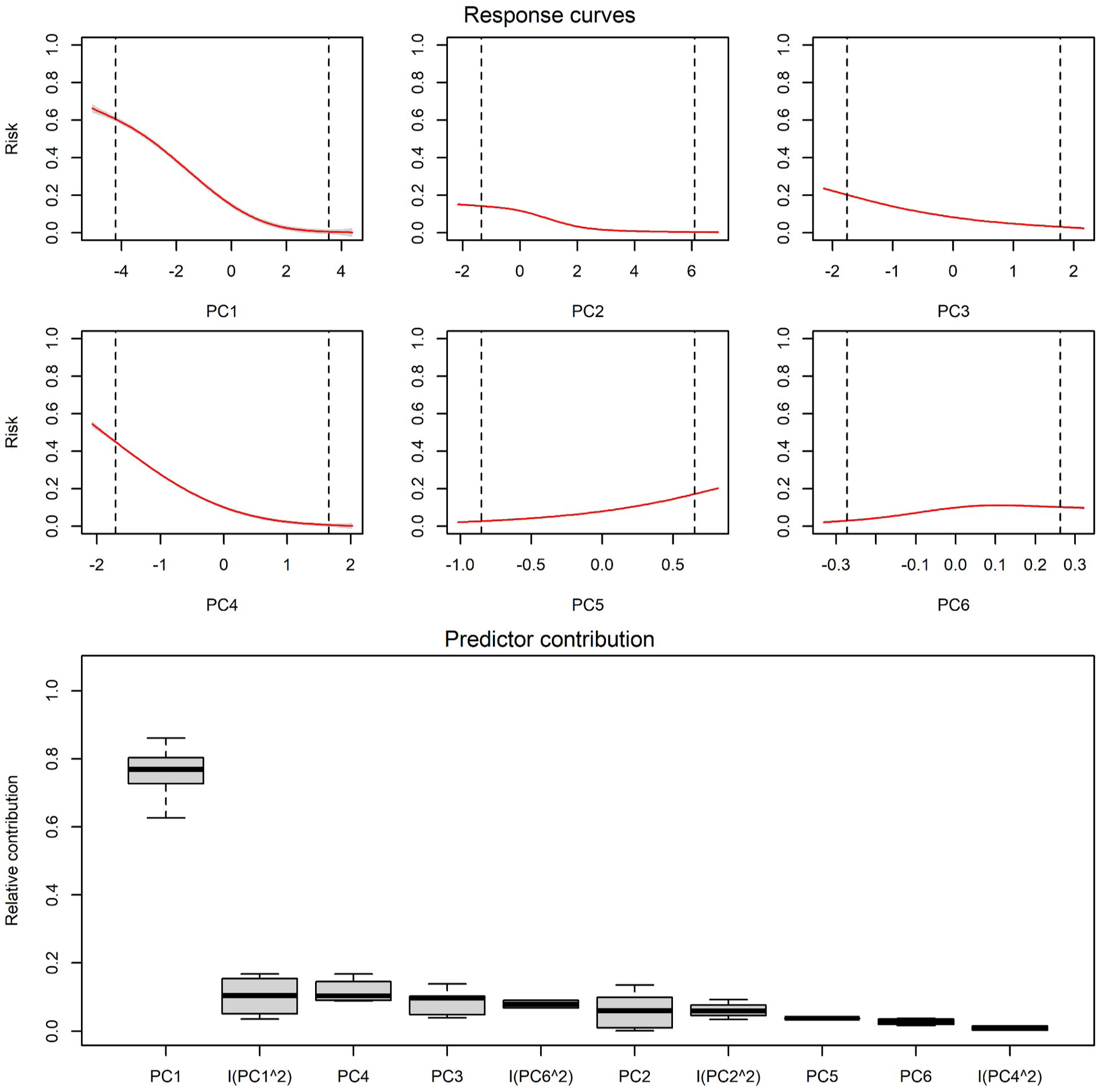
Ixodes scapularis

**Figures S5-S8.**
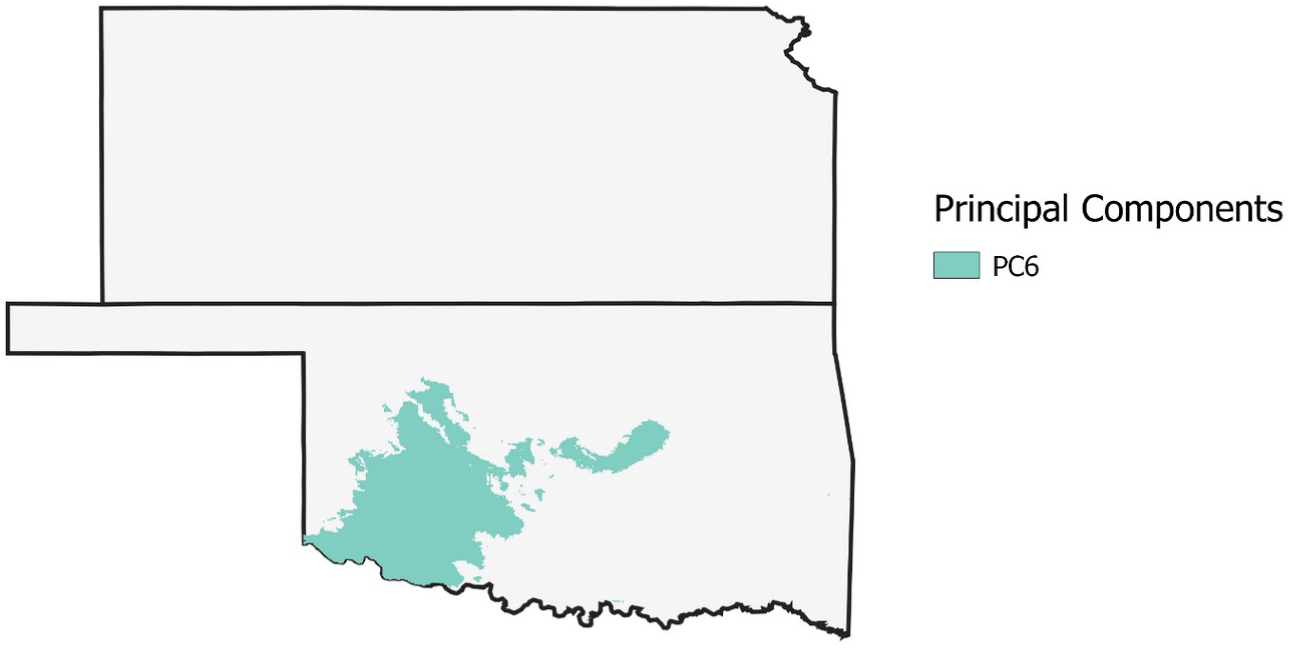
Contribution of different principal component axes to extrapolative areas in Kansas and Oklahoma in the MOP analysis. **S5.** January high:

**S6.**
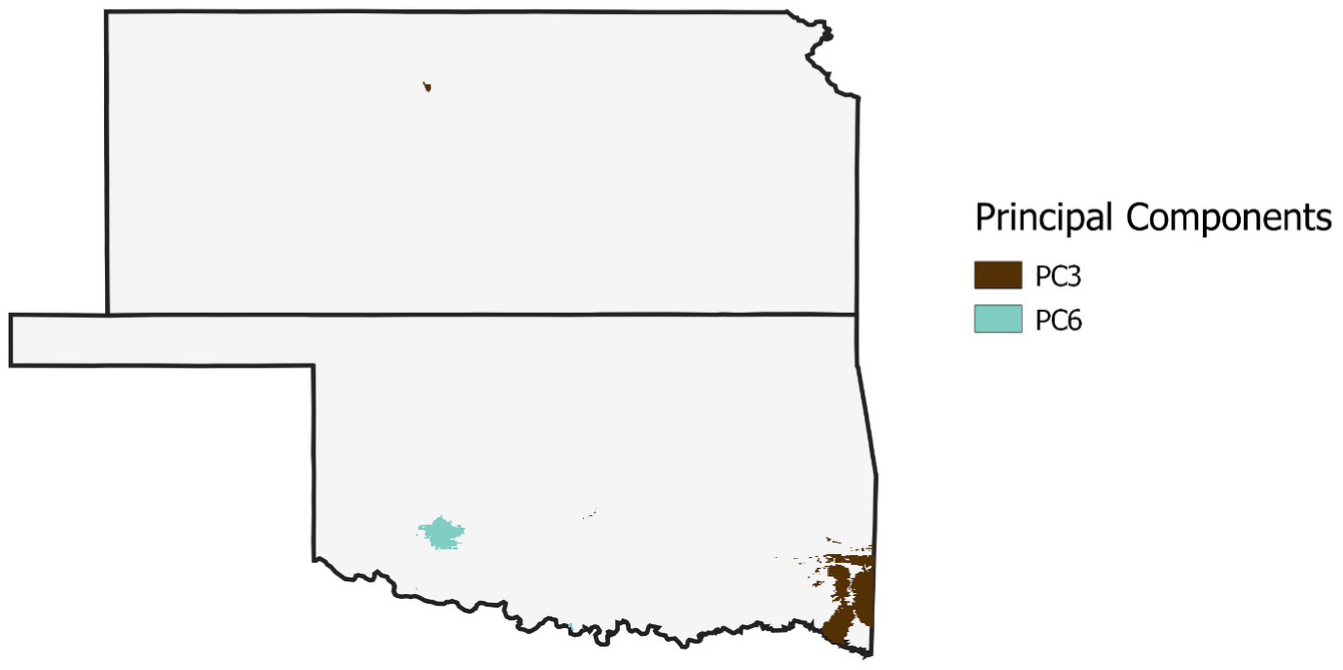
March high:

**S7.**
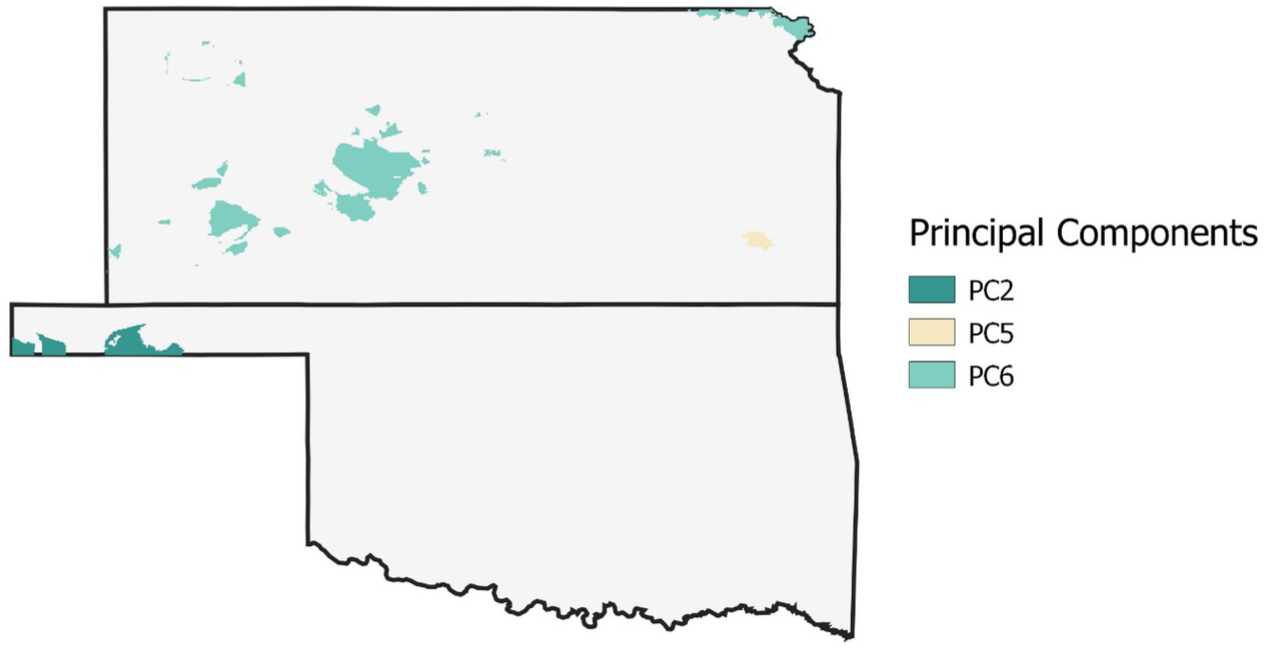
June low

**S8.**
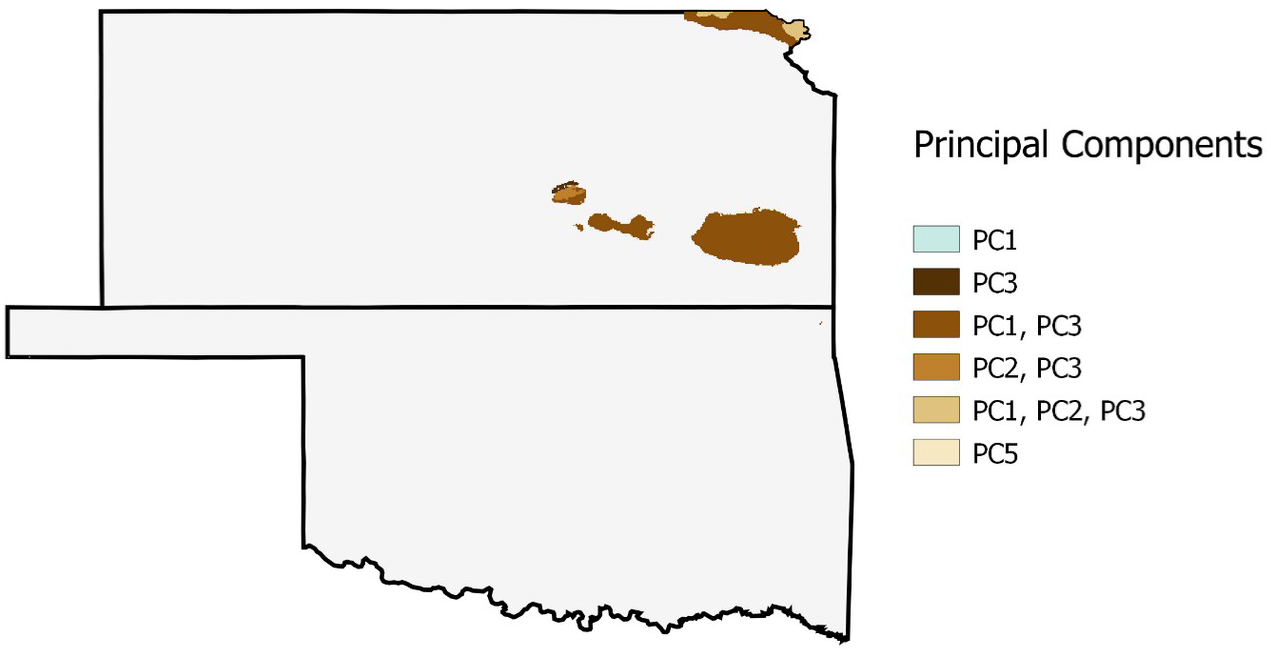
June high:

